# Resolved Genomes of Wastewater ESBL-Producing *Escherichia coli* and Metagenomic Analysis of Source Wastewater Samples

**DOI:** 10.1101/2024.03.12.584675

**Authors:** Clinton Cheney, Jared D. Johnson, John P. Ste. Marie, Kayla Y.M. Gacosta, Natalie B. Denlinger Drumm, Gerrad Jones, Joy Waite-Cusic, Tala Navab-Daneshmand

## Abstract

Extended-spectrum beta-lactamase (ESBL) producing *Escherichia coli* pose a serious threat to human health because of their resistance to the most commonly prescribed antibiotics: penicillins and cephalosporins. In this study, we provide a genomic and metagenomic context for the determinant ESBL genes of *E. coli* isolated from various wastewater treatment utilities in Oregon, USA. Class A beta-lactamase genes on chromosomes (*bla*CTX-M, *bla*TEM) were clustered with antibiotic resistance genes associated with other classes of antibiotics (sulfonamides and aminoglycosides) along with insertional elements. ESBL genes such as *bla*CTX-M, *bla*TEM, and *bla*SHV were also detected on conjugable plasmids of IncF and IncI incompatibility types. One novel IncF plasmid (pSHV2A_ESBLF) was identified in which carried a multi-drug resistance genotype (*bla*SHV-2A, *aadA22, aac(3), aph(6)*, *tetA*, and *sul1*) in addition to a *mer* (mercury resistance) operon, colicin, and aerobactin genes. Shotgun metagenomic analysis of the *E. coli*-originating wastewater samples showed the presence of class A beta-lactamases; however, the ESBL genes identified in the *E. coli* genomes were below the detection limits. Other ESBL-associated genes (*i.e.*, *bla*OXA.11, *bla*FOX.7, and *bla*GES.17) were identified in the wastewater samples and their occurrences were correlated with the core microbial genera (e.g., *Paraprevotella*). In both the *E. coli* genomes and the wastewater samples, tetracycline, aminoglycoside, and beta-lactam resistance determinants frequently co-occurred. The unique combination of whole-genome and metagenomic analysis provides a holistic description of ESBL-producing organisms and genes in the Oregonian wastewater system.

## INTRODUCTION

Antimicrobial-resistant (AMR) pathogens – referred to as the Silent Pandemic – are a major public health concern (1). In 2019, 4.95 million deaths were attributed to AMR pathogens globally, with the virulent *Escherichia coli* as the leading cause of these deaths (2). The U.S. Centers for Disease Control and Prevention has distinguished extended-spectrum beta-lactamase (ESBL)-producing Enterobacterales as one of the most serious threats facing humanity’s efforts against AMR (3). ESBL-associated genes confer resistance to a broad spectrum of the most commonly prescribed antibiotic class: beta-lactams, including penicillins and 1^st^ to 3^rd^ generation cephalosporins (4, 5).

ESBL-producing *E. coli* may be associated with multi-drug resistance (MDR), a classification of resistance to three or more classes of antibiotics (6). Moreover, antibiotic resistance genes (ARGs), including ESBL genes, can be carried on plasmids, which play a major role in the dissemination of antibiotic resistance in the environment via the horizontal gene transfer (HGT) mechanism of conjugation (6–8). IncF plasmids, which have a narrow host range of Enterobacteriaceae and are common within *E. coli* genomes, have been documented to carry *bla*CTX-M-type ESBL genes (9, 10). ESBL-associated IncF plasmids have also been described as MDR, either carrying determinants for cross-resistant efflux pumps (*i.e.*, *mac*B, *Emr*B, *Mdt*K) or harboring multiple classes of ARGs, providing the hosting *E. coli* strain with resistance to multiple antibiotics, such as aminoglycosides, macrolides, and tetracyclines (11, 12). Prevalence of ESBL-associated genes on plasmids alongside resistance to other antibiotic classes as well as antimicrobials such as metals informs the risks associated with the potential transfer of these plasmids from the host bacteria to other microbial communities.

ESBL-encoding genes are common in nature, having been found in bacterial isolates and environmental samples on all seven continents, including remote areas in Antarctica (13–19). Municipal wastewater and wastewater treatment utilities are important reservoirs for the prevalence and dissemination of ARGs. The ESBL genes within the class-A beta-lactamase family (*bla*CTX-M, *bla*TEM, and *bla*SHV) are commonly detected in wastewater according to a recent review (20). A 2019 survey of ESBL-producing *E. coli* isolated from wastewater in the U.S. showed a concerning rate of resistant phenotypes to third-generation cephalosporins, carrying *bla*TEM ESBL genes, as well as the carbapenemase *bla*VIM (21). Co-localization of virulence factors and ESBLs on the same plasmid have been reported in isolates from wastewater samples (22, 23). This co-localization is a concern because of the association between pathogenicity and limited clinical treatment options. With a highly diverse set of bacteria in biological processes such as activated sludge, the transfer of ARGs between a variety of bacteria could support the proliferation of AMR in effluent water and biosolids streams (24, 25). Given the serious threat of ESBL-producing *E. coli* and the link between wastewater and public health, further characterization of wastewater bacteria and the resistome associated with wastewater streams is necessary to inform risk assessment and subsequent policymaking.

In this paper, we resolved the genome of ESBL-producing *E. coli* isolates collected from wastewater utilities in Oregon whose AMR genotypes and phenotypes were previously characterized in our lab (26). Using a hybrid sequencing and assembly strategy (short- and long-read sequencing), we identified ARGs and virulence factors harbored on plasmids and chromosomes. We further characterized incompatibility types and the co-occurrences of ARGs and virulence factors on plasmids. We investigated the transferability of these plasmid-mediated ESBLs via conjugation. Finally, using shotgun metagenomic analysis of the *E. coli*-originated wastewater samples, we described the microbial community, the resistome composition, and the potential associations with plasmid-mediated ESBL-genes as well as other ARGs.

## METHODS

### *E. coli* Isolates and Wastewater Samples

*E. coli* strains were previously isolated from influent, secondary effluent, final effluent, and biosolids streams of eight wastewater treatment utilities across Oregon as described in our previous study (Table S1) (26). Wastewater samples were collected between January 2019 and September 2020. For liquid samples (*i.e.*, influent, secondary effluent, and final effluent), incremental volumes were filter-concentrated through a 0.45 µm mixed-cellulose ester membrane (Whatman, Kent, UK) until the filter clogged. The filtered volumes were up to 40 mL for influent and up to 400 mL for secondary and final effluents. The filter paper was fixed with 1 mL of 50% ethanol and stored at −20 °C until further processing. For biosolids, between 0.25 to 0.50 g of biosolids were transferred to sterile microcentrifuge tubes and stored at −20 °C until further processing.

### Whole Genome Sequencing and Analysis

Frozen stock cultures of *E. coli* isolated from the wastewater samples were streaked onto tryptic soy agar (TSA; Hardy Diagnostics, Santa Maria, CA) and grown for 24 hours at 37 °C before being transferred to tryptic soy broth (TSB, Hardy Diagnostics, Santa Maria, CA) and cultured under the same conditions. DNA was extracted from TSB cultures following the manufacturer’s instructions using the DNeasy Blood and Tissue kit (Qiagen, Carlsbad, CA). Purified DNA was quantified, and quality checked using the Qubit 4 (Invitrogen, Carslbad, CA) and NanoDrop™ One^©^ Micro UV-VIS Spectrophotometer (ThermoFisher Scientific, Waltham, MA). Long-read sequencing libraries were prepared using the Rapid Barcoding Sequencing kit (SQK-RBK004) and sequenced on a MinION system (Oxford Nanopore, Oxford, UK) following the manufacturer’s protocol. Short-read sequencing was performed using the Illumina MiSeq as described previously (26).

Hybrid assembly of MinION and Illumina reads was performed with Trycycler v0.5.0 and Flye v2.9 for each isolate (27, 28). Chromosomes were defined as circularized contigs with lengths of 4.6-5.0 Mbp, and plasmids were defined as circularized extrachromosomal contigs. BLAST alignment analysis was performed and visualized with BRIG (29). Chromosome and plasmid contigs were annotated with Prokka v1.14.5 for general annotation of coding sequences and NCBI’s AMRFinderPlus v3.10 and the CGE’s VirulenceFinder v2.0 were used to identify ARGs and virulence genes, respectively (30–32). Plasmid contigs were submitted to the pMLST web tool to determine associations with plasmid incompatibility groups (33).

### Conjugation Assays

The nine resolved ESBL-positive *E. coli* isolates (with the circularization of chromosomes and plasmids) in the study were used as donor strains in broth-mating conjugation assays with sodium azide-resistant *E. coli* J53 (ATCC BAA-2731, ATCC, Manassas, VA) as the recipient strain. Overnight cultures were grown at 37 °C in 50 mL Luria Broth (LB) supplemented with 5 mg/mL of cefotaxime (CTX) for donors and 100 mg/mL of sodium azide for the recipient. Overnight cultures were centrifuged at 4,500 × *g* for 15 minutes and resuspended in phosphate-buffered saline (PBS) twice to remove excess antibiotics. Donor and recipient cultures were mixed in equal proportions to a final volume of 1 mL before co-incubation for 1 hour at 22 °C. After incubation, mixtures were serially diluted in PBS and 70 µL of dilutions were spread on MacConkey agar with MUG plates (Hardy Diagnostics, Santa Clara, CA) containing 5 mg/mL CTX and 100 μg/mL sodium azide. The plates were incubated overnight at 37 °C to isolate transconjugant colonies. Six presumptive transconjugant colonies were randomly selected and grown overnight in 1 mL of LB prior to DNA extraction using the Wizard® Genomic DNA Purification kit (Promega, Madison, WI). The AMR genotypes of transconjugant colonies were determined by polymerase chain reaction (PCR) assays on a T100™ Thermal Cycler (Bio-Rad, Hercules, CA). Reaction volumes were 25 µL and consisted of 12.5 µL of Accustart II PCR Toughmix (Quantabio, Beverly, MA) and 0.2 µM concentration of forward and reverse primers. PCR assays involved an initial denaturation at 3 min at 94 °C, followed by amplification cycles consisting of 15 s of denaturation at 94 °C, 30 s of annealing, and 30 s extension at 72 °C for 35 cycles, followed by final extension of 7 min at 72 °C. Details of primers used in the study and associated annealing temperature are shown in Table S2. PCR products were viewed using a Gel Doc EZ Imager with Image Lab 5.2.1 software (Bio-Rad, Hercules, CA) on 3% agarose gel (VWR, Radnor, PA) with RedSafe Nucleic Acid Staining Solution (Bulldog Bio, Portsmouth, NH) and electrophoresis settings of 85 V for 45 minutes.

### Metagenomic Sequencing and Bioinformatics

DNA was extracted from wastewater samples using the FastDNA™ SPIN kit for Soil (MP Biomedicals, Irvine, CA) following manufacturer protocols and procedures described previously (34). For liquid samples (*i.e.*, influent, secondary effluent, and final effluent), the stored filter paper was removed from the ethanol solution, torn into small pieces, and transferred to the lysing tube. The remaining ethanol was centrifuged at 5,000 × *g* for 10 min and the supernatant was discarded. The pelleted cells were resuspended in the kit-supplied sodium phosphate buffer and transferred to the lysing tube. The remainder of the extraction process followed the manufacturer’s protocol. For biosolids, the manufacturer’s protocols were followed. DNA concentration and purity were determined using the NanoDrop™ One^©^. The identifiers (IDs) assigned to the wastewater samples (source IDs) in the metagenomics analysis section correspond to the same letters as those of the *E. coli* isolates associated with those samples (Table S1).

DNA samples were sequenced at the Center for Quantitative Life Sciences at Oregon State University (Corvallis, OR) on an Illumina HiSeq platform using a Nextera XT library preparation kit (Illumina, San Diego, CA). Primer and adaptor removal and read-trimming were performed with fastP using default settings for Nextera adapters (35). Trimmed reads were assembled with MEGAHIT v1.2.9 and taxonomic annotation was performed with Kaiju v1.8.2 (36, 37). Annotation of ARGs in the samples was performed using the ARGs-OAP Pipeline v2.0 (38). Alpha diversity, Bray-Curtis dissimilarity, and Spearman correlation statistical analyses for both microbial and ARG datasets were done using the R package vegan (39). Statistically correlated (*p* < 0.01) ARGs and microbial genera were visualized as a network with the R package igraph (40).

## RESULTS

### Distribution of ARGs in ESBL-producing *E. coli* genomes

Using the hybrid assembly of MinION and Illumina reads, nine of the eleven *E. coli* genomes were fully resolved with the circularization of chromosomes and plasmids. Two of the isolates, ESBL-A and ESBL-H, were not resolved. Of the nine resolved genomes, isolate ESBL-E had the longest chromosome, with a length of 5.04 Mbp, and isolate ESBL-I had the shortest chromosome with a length of 4.62 Mbp. In the nine resolved genomes, fifteen plasmids were identified and typed via pMLST, and six of these plasmids carried ESBL-associated genes (pTEM1 in ESBL-B, Fig. S1; pCTXM55 in ESBL-C, Fig. S2; pSHV2A in ESBL-F, Fig. 3; pTEM1 in ESBL-G, Fig. S3; pCTXM55 plasmid in ESBL-I *E. coli* isolate in ESBL-I, Fig. S4; and pCTXM55 in ESBL-J in Fig. S5). There were 83 ARGs identified by AMRFinder across the nine resolved genomes, with 63.9% (*n* = 53) of these ARGs located on the chromosomes and 36.1% (*n* = 30) contained on plasmids of *E. coli* isolates ESBL-B, ESBL-C, ESBL-F, ESBL-G, ESBL-I, and ESBL-J (Fig. 1, Table S3).

**FIG 1.**
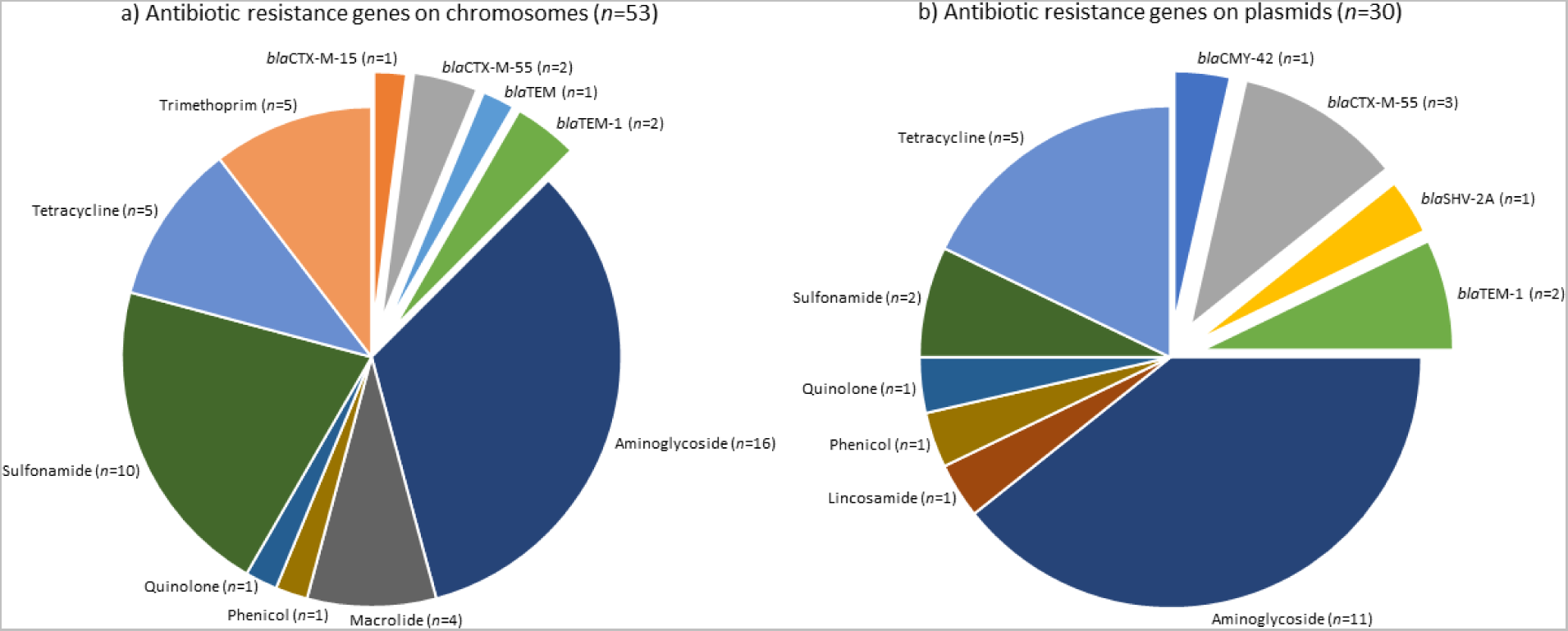
Distribution of antibiotic resistance genes on (a) chromosomes (*n* = 53) and (b) plasmids (*n* = 30) within the nine resolved *E. coli* genomes. The exploded slices in each pie demonstrate the ESBL-associated genes. The numbers in parenthesis show the incidences of these genes across the nine resolved *E. coli* genomes.

Aminoglycoside-resistance genes (*aac*3, *aad*A1, *aad*A5, *aad*A22, *aph*3, *aph*6) were the most common ARGs on both plasmids and chromosomes, making up 30.2% (*n* = 16) and 36.7 % (*n* = 11) of ARG presence, respectively (Fig. 1 and Table S3). ESBL genes were the second most abundant class of ARGs at 23.3% (*n* = 7) on plasmids (Fig. 1B), and at 11.3% (*n* = 6) on the chromosome (Fig. 1a). Five different ESBL genes were carried amongst the nine isolates: *bla*CTX-M-15, *bla*CTX-M-55, *bla*TEM-1, *bla*CMY-42, and *bla*SHV-2A. Of these ESBL genes, *bla*CTX-M and *bla*TEM types were the most common. *bla*CTX-M-15 was only seen on the chromosome of isolate ESBL-K. *blaCTX*-M-55 gene, however, was harbored by the chromosomes of ESBL-B and ESBL-D and plasmids of *E. coli* isolates ESBL-C, ESBL-I, and ESBL-J. *bla*TEM-1 was carried on plasmids of isolates ESBL-B and ESBL-G and chromosomes of ESBL-G, ESBL-I, and ESBL-J. The *bla*CMY-42 and *bla*SHV-2A genes were unique to their two hosts, ESBL-G and ESBL-F, respectively, and were both located on plasmids. Sulfonamide resistance genes (*sul*1, *sul*2) were relatively common on the chromosomes with six of the isolates carrying at least one, while only plasmids of isolates ESBL-F and ESBL-J had a *sul*1 and a s*ul*3 gene, respectively. The macrolide (*mph*A) and trimethoprim (*dfra*17) resistance genes were only chromosomally located in this isolate set. Tetracycline efflux transporters (*tet*A and *tet*B) were observed on five chromosomes and four plasmids.

### ESBLs and other ARGs clustered on *E. coli* chromosomes

The alignment and comparison of the nine resolved ESBL *E. coli* chromosomes are shown in Fig. 2. Seven of the nine resolved *E. coli* chromosomes carried ARGs as detected by AMRFinder, including six chromosomes that carried either *bla*CTX-M or *bla*TEM type ESBLs. BLAST alignment of the *E. coli* chromosomes shows over 90% nucleotide identity across the nine isolates, with three areas of dissimilarity where AMR genes were found in multiple isolates indicative of regions where accessory genes, such as ARGs, are located (Fig. 2). The majority of the annotated ARGs were clustered in these three variable regions and, interestingly, the ARGs were clustered together on their respective chromosomes with transposases including Tn*2*, Tn*As1*, Tn*As3*, IS*1A*, IS*1R*, IS*15*, IS*26*, IS*6100*, IS*Ecp1*, IS*Ec38*, IS*Vsa5*. The exception to the clustering of ARGs on these variable regions is the *bla*CTX-M-55 of Isolate ESBL-B which was approximately 2 Mbp apart from an ARG cluster (*i.e.*, *mph*A, *sul*1, *qac*E, *aad*A5, *dfra*A12, and *tet*A). In Isolates ESBL-C, ESBL-I, and ESBL-J, the chromosomal ARGs were all clustered in the same region between 2.67 and 2.68 Mbp (Fig. 2); all three of these isolates were identified as sequence type ST 744 (Table S1). It is notable that these three isolates with the same sequence type carried a similar array of ARGs in the same region of the chromosome, with a *sul*1*-qac*E1*-aad*A5*-dfr*A17*-*IS*26* region in full alignment with each other (Fig. 2). Isolate ESBL-G shared this region, but on the opposite strand and at a distinct region of the chromosome. Isolates ESBL-J and ESBL-I carried a *bla*TEM-1 followed by macrolide resistance *mph*A on the opposite strand, while isolate ESBL-C did not have an ESBL gene in this region nor anywhere else on its chromosome. As expected, all nine resolved chromosomes carried the *amp*C encoding beta-lactamase which was identical across all isolates and located in the identical region of the chromosome, at the end of a fumarase reductase operon (*fum*ABCD) and before a *blc* outer membrane lipoprotein (Fig. 2).

**FIG 2.**
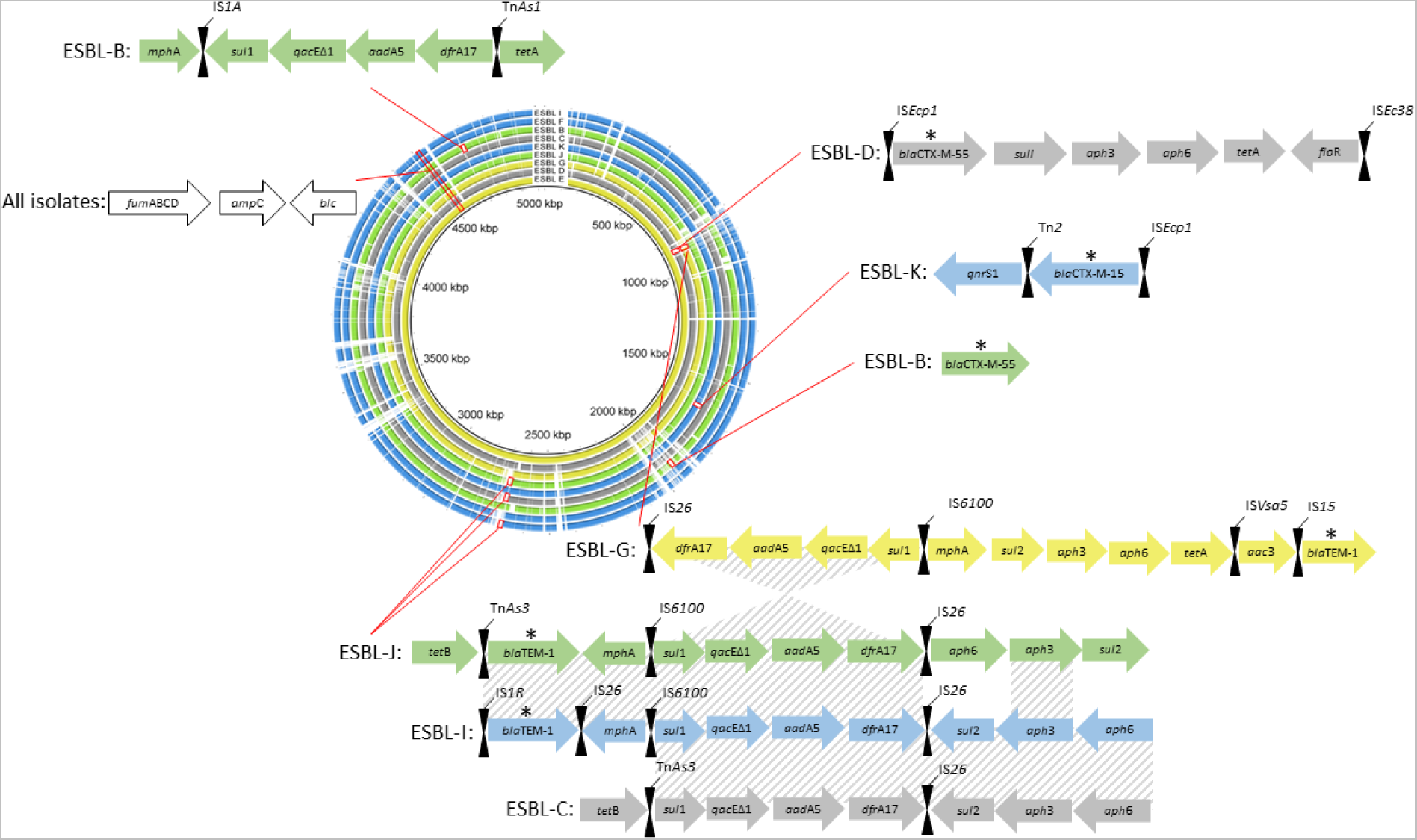
Alignment of the nine resolved ESBL-positive *E. coli* chromosomes and locations of their ARGs (arrows) and nearby transposases (hourglasses). Isolates are ordered by their chromosomes’ length from the longest in the innermost ring (isolate ESBL-E with a length of 5036705 bp) to the shortest in the outermost ring (isolate ESBL-I with a length of 4626801 bp) with the origin at the replication origin. Ring and annotation colors are according to the source sample type: green – influent, yellow – secondary effluent, blue – final effluent, and grey – biosolids. Areas of conserved synteny in the ARG regions are highlighted with diagonal stripes. ESBL genes are indicated by an asterisk over the arrows.

### Conjugable plasmids and their resistance determinants

In conjugation assays, all resolved isolates were mated with azide-resistant *E. coli* J53. Five of the nine isolates (ESBL-B, ESBL-C, ESBL-F, ESBL-G, and ESBL-J) produced transconjugants resistant to cefotaxime (100 µg/mL) and azide (5 µg/mL). These five isolates that produced transconjugants carried ESBL-associated genes on plasmids as observed in their resolved genomes (Table S3; extrachromosomal, circularized contigs). ESBL-I was the only isolate with a plasmid-borne ESBL gene that did not produce transconjugants in the assay. The antibiotic resistance genotypes of six randomly selected transconjugant colonies were tested via PCR. Generally, ARGs co-located with the ESBL genes on the same plasmids were observed in the transconjugants, while ARGs in the chromosomes of the donor isolates were not detected, demonstrating the transfer of the plasmids to the recipient *E. coli* (Table S4). The exception observed was for isolate ESBL-C where *sul*1 and *aad*A5 were detected in the transconjugants, whereas these genes were not assembled on ESBL-C’s plasmid (Fig. S2). Closer analysis showed that *sul*1 and *aad*A5 in ESBL-C were co-located between two identical terminal repeat regions on the chromosome which may have contributed to the misassembly of this region.

The ESBL-harboring plasmids of ESBL-C, ESBL-F, and ESBL-J were classified as IncF type, and were labeled pCTXM55_ESBLC, pSHV2A_ESBLF, and pCTXM55_ESBLJ, respectively (Figs. S2, 3, and S5). Alignment analysis of these three IncF plasmids and the closest BLAST hits to pSHV2A_ESBLF (which was the most unique plasmid identified in our study) is shown in Fig. 3. All three plasmids (pCTXM55_ESBLC, pSHV2A_ESBLF, and pCTXM55_ESBLJ) contained a *tra* conjugation operon identical to two plasmids previously deposited on GenBank (p1 accession CP059932.1 and pMCR-PA accession CP29748.1), as well as *rep*A, *psi*B, *umu*C and *sop*A genes related to plasmid maintenance/replication. All three IncF plasmids also carried the *cbi*, *cma*, and *cmi* colicin virulence factors. The 113-kbp long plasmid pCTXM55_ESBLC was identical to regions of two other plasmids observed in GenBank (p1 accession CP059932.1 and pRHB02-C06 accession CP058075.1) and contained 7 virulence factors including the colicins, *iuc*ABCD, *hly*F, and *omp*T (Fig. S2). pCTXM55_ESBLC had antibiotic resistance genotypes of tetracycline (*tet*A) and ESBL (*bla*CTX-M-55). The 122-kbp plasmid pCTXM55_ESBLJ (Fig. S5) was nearly identical to the same GenBank plasmid as pCTXM55_ESBLC (p1 accession CP059932.1), except for three ~200 bp regions that contained IS*tB,* Tn*pB, and* Tn*pA26* transposases. pCTXM55_ESBLJ had a MDR genotype for beta-lactams (*bla*CTX-M-55), tetracyclines (*tet*A) and aminoglycosides (*aac*3, *aad*A1, *aad*A22), sulfonamides (*sul*3), and florfenicol (*flo*R). pSHV2A_ESBLF was the most unique plasmid observed in our study (Fig. 3). This plasmid shared the conjugation and partitioning machinery of the other two IncF plasmids (pCTXM55_ESBLC and pCTXM55_ESBLJ) but was unique in the variable region (Fig. 3, 45 kbp – 90 kbp) that stored the plasmid’s ARGs. This region also carried a *mer* mercury resistance operon. pSHV2A_ESBLF contained ARGs for four antibiotic classes: beta-lactams (*bla*SHV-2A), aminoglycosides (*aad*A22, *aac*3, *aph*6), tetracyclines (*tet*A),and sulfonamides (*sul*1). The conserved conjugation and replication region was identical to two plasmids within GenBank: p113.6k (accession CP063370.1) and pPSEO2 (accession CP011916.1).

**FIG 3.**
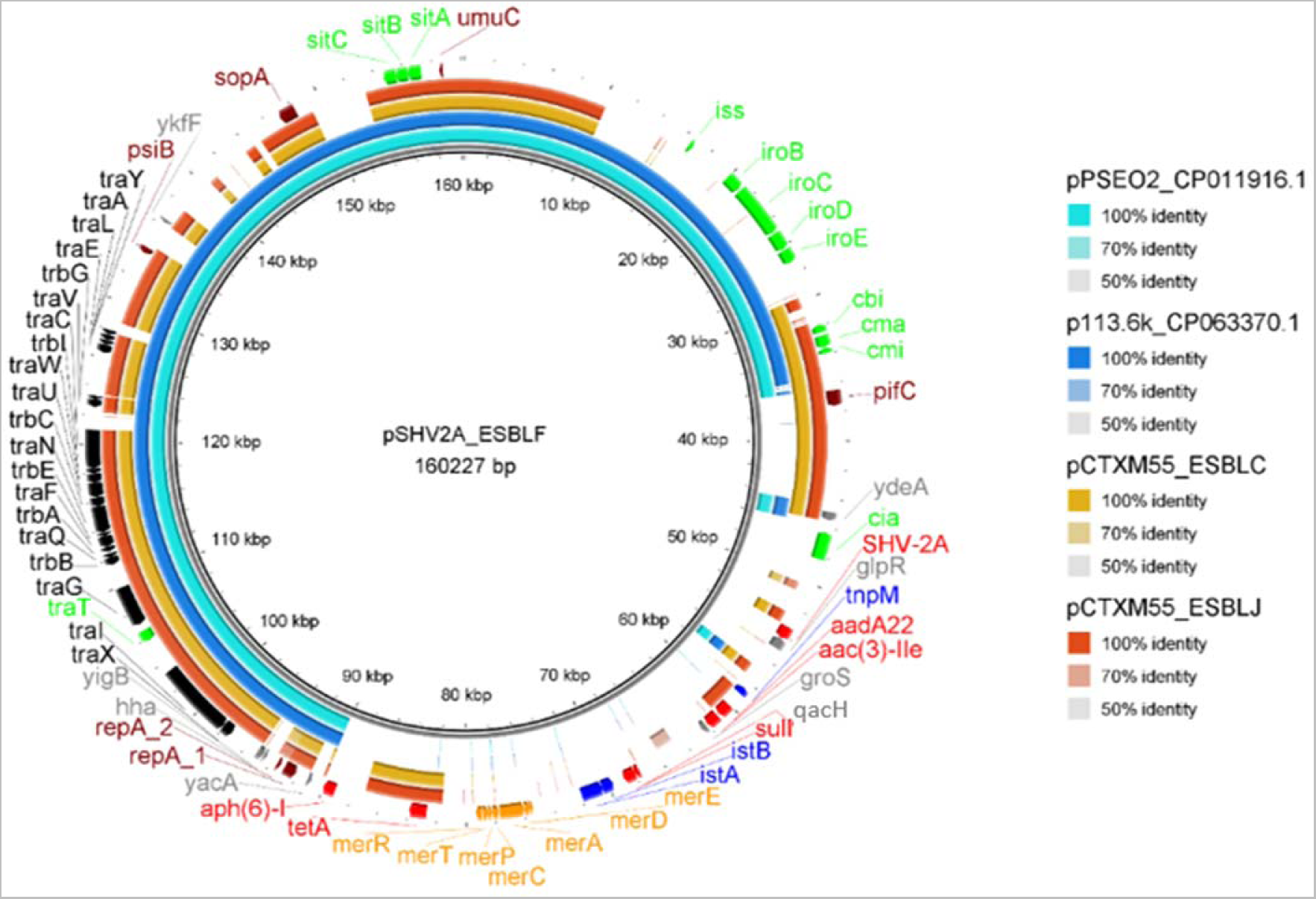
Alignment of pSHV2A plasmid in ESBL-F *E. coli* isolate and other similar ESBL-harboring IncF plasmids from this study as well as BLAST hits from the GenBank database. Outer two rings (dark and light orange) represent two similar IncF plasmids from this study (pCTXM55 plasmid in ESBL-J and pCTXM55 plasmid in ESBL-C *E. coli* isolates). Inner rings (dark and light blue) represent the closest matches to the pSHV2A plasmid in ESBL-F in the GenBank database. Gene labels are colored according to protein function: green – virulence factor, black – conjugation, maroon – plasmid replication/maintenance, red – antibiotic resistance gene, blue – mobile genetic element, and gold – mercury resistance.

Isolates ESBL-B and ESBL-G carried plasmids of the IncN and IncI incompatibility types respectively, named pTEM1_ESBLB (Fig. S1) and pTEM1_ESBLG (Fig. S3) with different conjugation systems than the IncF plasmids in Fig. 3. pTEM1_ESBLB, the smallest ESBL-harboring plasmid observed in these isolates (43.8 kbp), had a *pil*X operon related to conjugable pili generation and a *res*-*par*GF plasmid partitioning system. The *bla*TEM-1 ESBL gene was the only ARG on this plasmid, with similarity to the plasmid pOLA52 (accession EU370913.1). pTEM1_ESBLG, a 69.7 kbp IncI type plasmid, harbored a *bla*TEM-1 ESBL and *bla*CMY-42 beta-lactamase along with a *qnr*S1 quinolone resistance gene. Interestingly, *bla*CMY-42 was flanked by a *blc* lipoprotein, similar to the *amp*C gene detected in all nine resolved chromosomes. No virulence factors in the VirFinder database were detected on this plasmid. The area containing ARGs on pTEM1_ESBLG was not detected on any plasmids in GenBank, while the area containing the conjugation genes *tra*A*-trb*AB*-nik*AB matched those of the previously described plasmid pS68 (accession KU130396.1). The plasmid assembled from isolate ESBL-I (pCTXM55_ESBLI) was not observed to transfer via conjugation but carried *bla*CTX-M-55, *tet*A, *and aac*3 ARGs, aerobactyn synthetase virulence factors (*iuc*BCD), and IncI-type alleles. The one-hour conjugation test may not have been long enough to observe the transfer of this plasmid, possibly because the conjugation rate was inhibited via competition with another, non-ARG-carrying IncF plasmid observed in this genome.

### Virulence factors co-located with ESBLs

Within the nine resolved *E. coli* genomes, we identified a total of 115 virulence factors (Table 1). 67% (*n* = 77) of the virulence factors were located on chromosomes, with all nine chromosomes carrying at least one virulence factor. The other 38 virulence factors were located on plasmids that also harbored ESBL-encoding genes (Table 1). Notable were the colicin virulence factors (*cba, cia, cib,* and *cma*) with at least one observed on all six resolved plasmids that carried ESBL genes. Aerobactin synthetase and receptor genes *iuc*C and *iut*A were observed on the three plasmids that carried *bla*CTX-M-15. Colicin virulence factors, *iuc*C, and *iut*A were all unique to plasmids, with no occurrences on chromosomes. In contrast, the virulence-related iron transport protein *sit*A was found in three chromosomes (ESBL-B, ESBL-E, and ESBL-K), and in four plasmids (ESBL-C, ESBL-F, ESBL-I, ESBL-K). The most common virulence factors in their genomes, glutamate decarboxylase (*gad*) and tellurium ion resistance protein *ter*C, were chromosomal only, with multiple isolates having redundant copies of these genes.

**TABLE 1.** Distribution of virulence factors in ESBL-producing *E. coli* isolates.

| Virulence factor function | Gene(s) | ESBL *E. coli* isolate ID |  |  |  |  |  |  |  |  |
| --- | --- | --- | --- | --- | --- | --- | --- | --- | --- | --- |
|  |  | B | C | D | E | F | G | I | J | K |
| Aerobactin synthetase | *iucC* |  | P<sup>1</sup> |  | – |  |  | P<sup>1</sup> | P<sup>1</sup> |  |
| Afimbrial adhesion | *afaD* |  |  |  |  |  | C |  |  |  |
| Capsule polysaccharide export | *kpsE* |  |  | C | C |  |  |  |  |  |
| Colicin | *cba, ci(ab), cma* |  | P<sup>2</sup> |  | P<sup>2</sup> | P<sup>2</sup> |  | P<sup>2</sup> | P<sup>2</sup> | P<sup>2</sup> |
| Enterobactin siderophore receptor protein | *iroN* |  |  |  |  | P<sup>3</sup> |  |  |  |  |
| Ferric aerobactin receptor | *iutA* |  | P<sup>1</sup> |  |  |  |  | P<sup>1</sup> | P<sup>1</sup> |  |
| Glutamate decarboxylase | *gad* | C | C | C | C | C | C | C | C | C |
| Heat-resistant agglutinin | *hra* |  |  | C |  |  |  |  | C |  |
| Hemolysin F | *hlyF* |  | P<sup>1</sup> |  |  | P<sup>3</sup> |  | P<sup>1</sup> | P<sup>1</sup> |  |
| High molecular weight protein 2 non-ribosomal peptide synthetase | *irp2* |  |  |  | C |  |  |  |  |  |
| Increased serum survival | *iss* | C |  | C |  | C, P<sup>3</sup> |  |  |  |  |
| Iron transport protein | *sitA* | C | P<sup>1</sup> |  | C | P<sup>3</sup> |  | P<sup>1</sup> | P<sup>1</sup> | C |
| Long polar fimbriae | *ipfA* |  |  | C |  |  | D |  |  | D |
| Major pilin subunit F11 | *papA_F11* |  |  |  |  |  |  |  | C |  |
| Major pilin subunit F19 | *papA_F19* |  |  | C |  |  |  |  |  |  |
| Microcin | *cvaC, mcmA* |  | P<sup>1</sup> |  |  | P<sup>3</sup> |  | P<sup>1</sup> | C |  |
| Outer membrane hemin receptor | *chuA* |  |  | C | C |  |  |  |  |  |
| Outer membrane protease (protein protease 7) | *ompT* |  | P<sup>1</sup> |  | C | P<sup>3</sup> |  | P<sup>1</sup> | P<sup>1</sup> |  |
| Outer membrane protein complement resistance | *traT* |  | P<sup>1</sup> |  |  | P<sup>3</sup> |  | P<sup>1</sup> | P<sup>1</sup> |  |
| Outer membrane usher | *papC, afaC* |  | C |  |  |  |  |  | C |  |
| Periplasmic chaperone | *afaB* |  |  |  |  |  |  |  | C |  |
| Polysialic acid transport protein; Group 2 capsule | *kpsMII_K5* |  | C |  |  |  |  |  |  |  |
| Siderophore receptor | *fyuA, ireA* |  |  |  | C |  |  |  | C |  |
| Tellurium ion resistance protein | *terC* | C | C |  | C | C | C | C | C | C |
| Transcriptional regulator | *afaA* |  |  |  |  |  |  |  | C |  |
| EAST-1 heat-stable toxin | *astA* |  | C |  |  |  |  |  |  |  |
| Enteroaggregative immunoglobulin repeat protein | *air* |  | C |  | C |  |  |  |  |  |
| *Salmonella* HilA homolog | *eilA* |  | C |  | C |  |  |  |  |  |

294 <sup>1</sup> on *bla*CTX-carrying plasmid295 <sup>2</sup> on all ESBL-carrying plasmids296 <sup>3</sup> on *bla*SHV-carrying plasmid

### Metagenomics

The alpha diversity indices (*i.e.*, Shannon diversity, richness, and evenness) of the microbial genera and ARGs for the eleven wastewater samples are provided in Table S5. The number of reads aligned to the microbial genera ranged from 5000 to 5500 except for final effluent Sample I, which was an outlier with only 3220 of its reads aligned. This sample’s microbial composition had a Shannon index of 4.96, similar to those of the three influent samples A, B, and J (4.91, 4.77, and 4.93, respectively). All other samples had Shannon indices of 6.0 to 6.3 in the genera dataset. The resistomes of influent samples A, B, and J had richness that ranged between 355 and 378 unique ARGs, while the remainder of the samples were less rich, with 144 to 277 unique ARGs. Shannon diversity indices for ARG abundance ranged from 3.09 to 4.48 in all samples, indicating that all the elevn wastewater samples had similar diversity of unique ARGs.

Bray-Curtis dissimilarity scores (herein referred to as dissimilarity) and NMDS plots are provided for the microbial community and resistome datasets in Figs. S6 and S7a and b, respectively. Biosolids samples (C, D, H) were similar to each other with respect to their microbial community composition (dissimilarity = 0.196) and, to a lesser degree, their resistome (dissimilarity = 0.363). The same applied to secondary effluent samples (E, G) (dissimilarity = 0.288 for genera, 0.363 for ARGs). Influent samples (A, B, J) were equally similar with regard to both datasets (dissimilarity = 0.276 for genera, 0.222 for ARGs). Effluent samples (F, I, K) were the most dissimilar sample types to each other with dissimilarity scores greater than 0.5 in both datasets. PERMANOVA analysis identified that for both the microbial community and the resistome, the dissimilarities between each treatment group (*i.e.*, sample type) were significantly higher than the differences within each treatment group (microbial community: *p* < 0.001; resistome: *p* < 0.01).

The top microbial genera (>0.1% relative abundance) and ARGs (>0.02 gene copies/16S rRNA) are displayed in the heatmaps in Figs. 4a and b, respectively. *Candidatus cloacimonas* was the most abundant genus in all three of the biosolids samples (D, C, H), while sparser in other sample types, contributing to the clustering of these samples together in the NMDS plot in Fig. S7a. The genera *Acinetobacter* (11.0%), *Acidovorax* (15.0%), *Arcobacter* (5.8%), *Aeromonas* (5.4%), and *Bacteroides* (4.1%) were relatively abundant in influent samples and present in secondary effluent, and final effluent samples. *Flavobacterium*, *Pseudomonas*, *Rhodoferax,* and *Dechloromonas* were more abundant in secondary and final effluent samples compared to influent, likely because the total microbial count was lower in primary and secondary treatment, and these were the primary genera remaining. The bacitracin *bac*A, aminoglycoside *aad*A, sulfonamide *sul*1, and multidrug *acr*B were common ARGs in all eleven samples. The macrolide resistance gene *erm*F was the most common ARG in biosolids samples, while not as common in other sample types. The most prevalent ESBL genes in these wastewater samples were Type A beta-lactamases, which is the class that includes *bla*SHV, *bla*TEM, and *bla*CTX-M type ESBLs. These were the most abundant in the secondary effluent sample E and the final effluent sample K. The ESBL genes *bla*AER.1 and *bla*CfxA2 were also detected in the same samples as class A beta-lactamases. Species abundance data was parsed for ESKAPE pathogens (six highly virulent and antibiotic-resistant bacterial pathogens including: *Enterococcus faecium*, *Staphylococcus aureus*, *Klebsiella pneumoniae*, *Acinetobacter baumannii*, *Pseudomonas aeruginosa*, and *Enterobacter spp*., as well as *E. coli*; Fig. 4c). The most abundant ESKAPE pathogen species in all sample types was *P. aeruginosa* which ranged from 0.6 to 2.7% of the microbial community (Fig. 4c). *S. aureus* and *E. faecium* were below 0.01% abundant in all eleven samples. *E. coli* was less than 0.1% of the microbial community in all samples.

**FIG 4.** Microbial community and resistome of ESBL-producing *E. coli*-originating wastewater samples. Letters A-K correspond to wastewater sample IDs. a) Heatmap of the relative abundance of the microbial genera present above 0.1% (log-scale). b) Heatmap of the anitbiotic resistance genes concentrations (gene copies/16S rRNA, log-scale) grouped by their corresponding antibiotic class. c) Relative abundance of ESKAPE pathogens and *E. coli* with samples grouped according to wastewater sample type.

Of the ARGs found in *E. coli* isolates, the most abundant in the *E. coli*-originating wastewater samples were *sul*1 (0.004 – 0.023 gene copies/16S rRNA), *aad*A (0.002 – 0.014 gene copies/16S rRNA), *qac*EΔ1 (0.001 – 0.020 gene copies/16S rRNA), and *tet*A (0.001 – 0.005 gene copies/16S rRNA) (Table S6). The ESBL genes *bla*CTX-M-55 and *bla*TEM-1, which were identified in *E. coli* isolates, were detected in one (Sample B) and five wastewater samples (Samples B, E, F, J, and K) from five unique facilities, respectively, at levels greater than 0.001 gene copies/16S rRNA. ESBL genes *bla*CTX-M-15, *bla*SHV-2A and *bla*CMY-42, which were also identified in *E. coli* isolates, were undetected in all source samples. Seven beta-lactam ARGs were detected above 0.001 gene copies/16S rRNA in at least three samples (Table S7): class A beta-lactamases (<0.001 – 0.061 gene copies/16S rRNA), *bla*AER-1 (<0.001 – 0.013 gene copies/16S rRNA), *bla*CfxA2 (<0.001 – 0.011 gene copies/16S rRNA), *bla*OXA-2 (<0.001 – 0.009 gene copies/16S rRNA), *bla*OXA-10 (<0.001 – 0.007 gene copies/16S rRNA), *bla*CfxA3 (<0.001 – 0.002 gene copies/16S rRNA), and *bla*OXA-119 (<0.001 – 0.002 gene copies/16S rRNA). None of the specific ESBL-genes found in the *E. coli* isolates were found to be significantly correlated (*p* < 0.01) with the microbial genera in *E. coli*-originated wastewater samples. However, correlation analysis demonstrates significant associations between the microbial genera with 39 other ESBL-associated genes (Fig. 5). The prevalence and abundance of *Cloacibacterium* genera, which was only detected above 0.01% abundance in all three influent samples (A, B, J) and one final effluent sample (F), correlated with *bla*OXA.58 and *bla*MOX.2,5,6 beta-lactamases (Spearman’s ρ = 0.989, *p* < 0.01). *Paraprevotella* correlated with *bla*OXA.11, *bla*FOX.7, *bla*GES.17, and *bla*OXA.164 (Spearman’s ρ = 0.985, *p* < 0.01), along with the colistin resistance gene *mcr.1.9* (Spearman’s ρ = 0.985, *p* < 0.01) and the multidrug efflux pump *sme*F (Spearman’s ρ = 0.985, *p* < 0.01). These were in the same cluster as the genera *Faecalibacterium* and *Roseburia* with the ESBL-associated gene *bla*TEM.187. Another colistin resistance gene *mcr.3* correlated with the class C beta-lactamase class (Spearman’s ρ = 0.986, *p* < 0.01), which is inclusive of *amp*C enzymes. The ESKAPE pathogen genus *Klebsiella* was correlated with *bla*MOX.1 (Spearman’s ρ = 1.000, *p* < 0.01) and the multidrug ARG *lmr*P (Spearman’s ρ = 1.000, *p* < 0.01). The ESBL-related genes *bla*SHV.1 and *bla*SHV.4 co-occurred with the tetracycline resistance gene *tet*Y (Spearman’s ρ = 0.985, *p* < 0.01) in final effluent, secondary effluent, and influent samples. The ESBLs *bla*TEM.6, *bla*SHV.152, and *bla*OXA.7 were also significantly correlated with each other (Spearman’s ρ = 0.985, *p* < 0.01).

**FIG 5.**
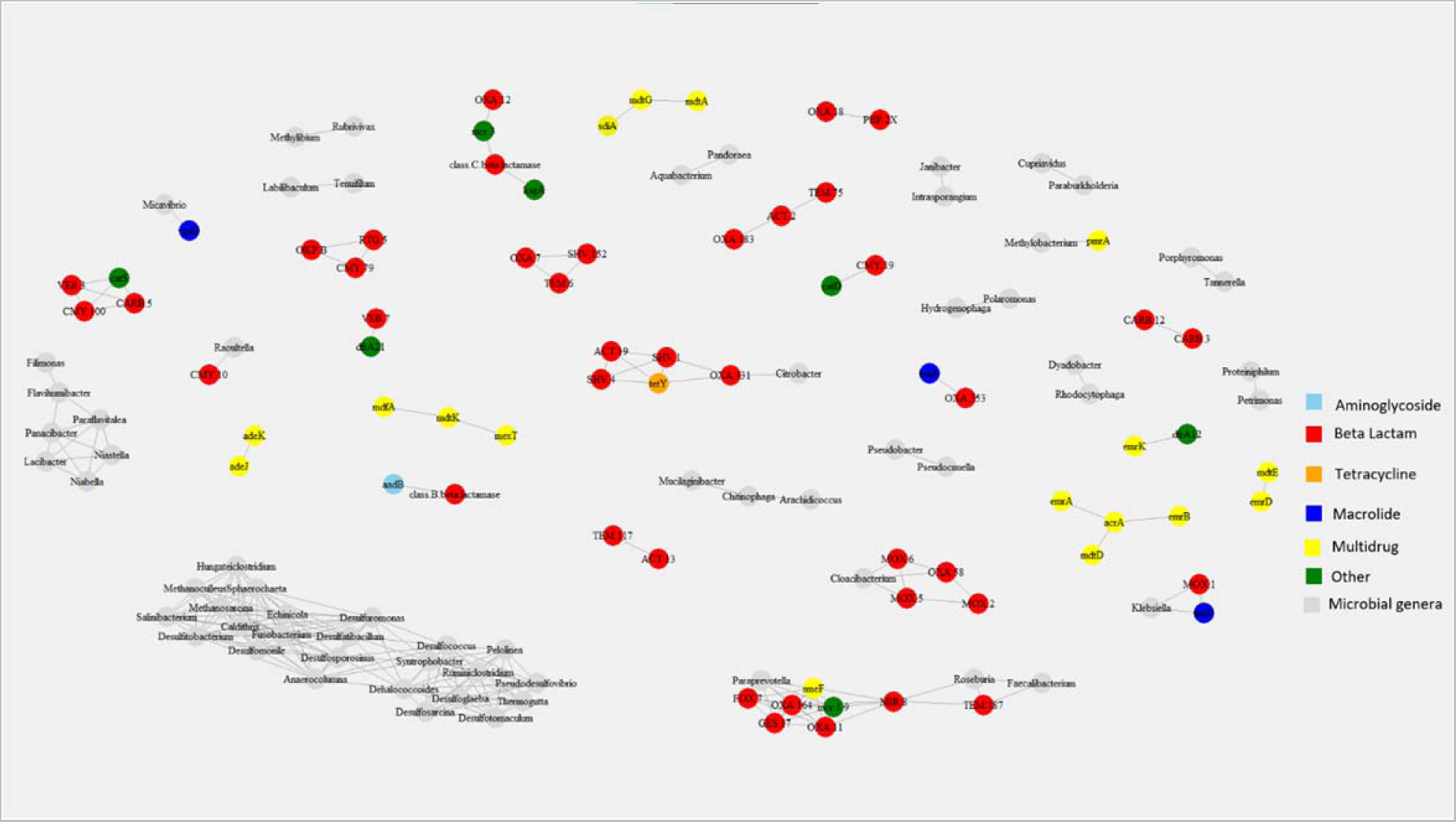
Network analysis of microbial genera and antibiotic resistance genes in wastewater samples. Edges represent significant Spearman correlation (*p* < 0.01) of nodes with a neighborhood of three or more. Colored symbols indicate antibiotic resistance genotypes and grey symbols represent the microbial genera.

## DISCUSSION

Worldwide, class A beta-lactamases, such as those derived from *bla*TEM, *bla*CTX, and *bla*SHV genes, are the most detected in Enterobacterales in the agricultural supply chain (41). Here, we detected class A beta-lactamases genes in several of the ESBL-producing *E. coli* and in their originating wastewater samples. These ESBL-producing *E. coli* isolates carried ARGs associated with nine antibiotic classes distributed on both chromosomes and plasmids. On chromosomes, ARGs were co-located together within variable regions in close proximity to various transposases or insertion elements (Fig. 2). This co-localization of mobile genetic elements and ARGs suggests that transposition events were originally responsible for the accumulation of these ARGs in the genome. Moreover, there was conserved synteny between the ARGs of three isolates (ESBL-C, ESBL-I, and ESBL-J, collected from three different samples from two different utilities, Table S1) belonging to sequence type ST 744 at the *sul*1*-qac*E1*-aad*A5*-dfr*A17*-*IS26 region, indicative of a similar origin for these ARGs in these three *E. coli* isolates. All nine *E. coli* isolates in this study were phenotypic ESBL-producers and carried a chromosomal *amp*C beta-lactamase that was homologous across the chromosomes. In combination with ESBL genes like *bla*TEM-1 and *bla*CTX-M-55, overexpression of these chromosomal *amp*C genes can have an additive effect on their beta-lactam resistance phenotype (42).

One aspect that makes the findings from this study unique and novel is the resolution of the genomes. Using a hybrid sequencing and assembly strategy (short- and long-read sequencing), we fully resolved the genomes of nine wastewater-originated *E. coli* isolates with the circularization of chromosomes and plasmids. Our method allowed us to identify the locations and orientations of ARGs and virulence factors harbored on plasmids and chromosomes of the nine ESBL-producing *E. coli* isolates.

The detection of ESBL-associated genes on conjugable plasmids indicates a potential for these genes to spread horizontally in wastewater treatment utilities and in downstream ecosystems (Table S3). The identification of transposases nearby ARGs on plasmids reflects the chromosomal ARG organization, for example, pCTX_ESBLJ carried an IS*26*-related element (Tn*pA26*) within a few hundred bp of aminoglycoside, sulfonamide, and tetracycline ARGs (Fig. S5). This may be indicative of the intercellular transposition of ARGs between chromosomes and plasmids. Five of the isolates were observed to transfer plasmid mediated ARGs between *E. coli* strains (Table S4). Three of these plasmids belonged to the IncF incompatibility group (Figs. S2, 3, and S5). These have been described as narrow-host range, and may only transfer to other Enterobacteriaceae or remain stable within *E. coli* genomes (9). A unique plasmid pSHV2A_ESBLF was resolved which carried a variable region with five classes of ARGs, a metal resistance operon (*mer*), and *iro*BCD and *sit*ABC virulence factors (Fig. 3). The co-localization of MDR determinants and metal resistance genes is concerning given the threat of antimicrobial-resistant pathogens in the natural and built environment (43).

Conjugation assays give valuable insight into the transferability of plasmids containing ARGs. However, the behavior of these organisms at the lab scale is likely not representative of the extent that horizontal gene transfer occurs within wastewater treatment systems. The narrow-host range plasmids that carry these ESBLs may lack suitable hosts in the wastewater treatment ecosystem, as our metagenomic analysis showed a low (< 0.1%) abundance of Enterobacterales. It should be noted that the conjugation assay used in this paper selects for specific transconjugants, with cefotaxime and azide phenotypes. The selective pressure to isolate these transconjugants may not exist in a wastewater system. Furthermore, the transfer of ESBL genes is not limited to plasmids; the ability for ARGs to be relocated from chromosomes to conjugatable plasmids through transposition is a contributing factor to the mobilization of ARGs (44).

Metagenomics showed that multidrug efflux pumps, bacitracin resistance genes, and class A beta-lactamases were among the most common ARGs across the eleven Oregon wastewater samples from which the ESBL-producing *E. coli* were isolated (Fig. 4b). Community analysis of the wastewater samples showed *Acinetobacter, Acidovorax, Aeromonas,* and *Bacteroides* as the core genera in influent, secondary effluent, and final effluent (Fig. 4a). *Candidatus cloacimonas* was a dominant genus in the three biosolids samples, all of which were from separate wastewater treatment utilities (Fig. 4a). A correlation analysis showed that the beta-lactam resistance genes *bla*TEM.8, *bla*SHV.152, and *bla*OXA.7 co-occurred at similar rates within wastewater samples. The ARGs *sul*1 and *aad*A occurred in both wastewater samples as well as the *E. coli* isolates. Presence of *E. coli* was low (< 0.1%) in these wastewater samples, but the location of these genes on conjugable plasmids and on IS*26* insertional elements in the *E. coli* genomes may explain the prevalence of these sulfonamide and aminoglycoside genes within the samples. While class A beta-lactamases, in general, were abundant (> 0.01 gene copies/16S rRNA), the specific ESBL genes *bla*CTX-M-55, *bla*CTX-M-15, *bla*TEM-1, *bla*SHV-2A, and *bla*CMY-42 that occurred in the *E. coli* isolates were below the detection limit in the *E. coli*-originating wastewater samples.

## CONCLUSION

Here we provide the genetic context of ESBL-producing *E. coli* isolated from wastewater. The association of ESBL genes and other classes of antibiotic resistance determinants with mobile genetic elements like IS*26* and conjugable plasmids indicates a horizontal transfer origin of these ARGs to the genomes. The clustering of ARGs together in specific regions of chromosomes indicates that these ARGs have become fully integrated into the *E. coli* genomes through previous horizontal gene transfer events. The occurrence of plasmids associated with MDR and virulence genotypes, including a novel plasmid that also carried a mercury resistance operon, is concerning given the role conjugation plays in the spread of antibiotic resistance and virulence in the environment. However, the ESBL-harboring plasmids in the *E. coli* genomes were all characterized as narrow-host range, so it is possible that these genes are relatively stable in their hosts. Additionally, the specific ESBL genes *bla*CTX-M-55, *bla*CTX-M-15, *bla*TEM-1, *bla*SHV-2A, and *bla*CMY-42 that occurred in the *E. coli* isolates were in low abundance (<0.001 gene copies/16S rRNA) in the *E. coli*-originating wastewater samples. This may be associated with the low relative abundance of Enterobacterales in wastewater samples. Another possibility is that beta-lactam resistance genes are diverse between different microbial genera. Furthermore, class A beta-lactamases, bacitracin, and multidrug efflux pumps were the most abundant types of ARGs found in the eleven wastewater samples. Correlation of core microbial genera of wastewater samples, such as *Cloacibacteria,* with ESBL-related genes, indicates that ESBLs are intrinsic to the Oregonian wastewater treatment system. The ESBL-associated genes *bla*OXA.11, *bla*FOX.7, and *bla*GES.17 were not found in any of the *E. coli* isolates but were associated with other wastewater microbial genera like *Paraprevotella, Faecalibacterium,* and *Roseburia*. The cross-genera spread of MDR plasmids would be valuable for future research efforts. The detection of a broad spectrum of ARGs and ESBL-producing *E. coli* isolates in biosolids and final effluents is concerning given that these streams are transported to downstream ecosystems like rivers and agricultural fields. This study’s unique incorporation of hybrid whole-genome sequencing, culture-dependent methods, and metagenomics provides a holistic picture of ESBL-producing *E. coli* within Oregonian wastewater systems, and by proxy Oregonian communities.

## ACKNOWLEDGMENTS

This work was supported by the USDA National Institute of Food and Agriculture, Agricultural and Food Research Initiative Competitive Program, Agriculture Economics and Rural Communities, Grant No. 2018-67017-27631, in-kind supplement from the Oregon State University’s Center for Quantitative Life Sciences, and Oregon State University’s Agricultural Research Foundation. The authors would like to thank Dr. Hussein M.H. Mohamed (Oregon State University) for his assistance with DNA extraction and nanopore sequencing of ESBL-producing isolates.

## DATA AVAILABILITY

The sequenced data sets have been deposited in NCBI: the *E. coli* genomes are available in GenBank under BioProject accession number PRJNA1044148 and the wastewater metagenomes are available in Short Read Archive (SRA) under BioProject accession number PRJNA1060321.

